# High fidelity estimates of spikes and subthreshold waveforms from 1-photon voltage imaging *in vivo*

**DOI:** 10.1101/2020.01.26.920256

**Authors:** Michael E. Xie, Yoav Adam, Linlin Z. Fan, Urs L. Böhm, Ian Kinsella, Ding Zhou, Liam Paninski, Adam E. Cohen

**Affiliations:** Department of Chemistry and Chemical Biology, Harvard University, Cambridge, MA, USA; Department of Statistics, Columbia University, New York, NY, USA; Howard Hughes Medical Institute, Chevy Chase, MD, USA

**Author notes:** Edmond and Lily Safra Center for Brain Sciences, Hebrew University, Jerusalem, 91904 Israel (Yoav Adam). Department of Bioengineering, Stanford University, Stanford, CA, USA.

## Abstract

The ability to probe the membrane potential of multiple genetically defined neurons simultaneously would have a profound impact on neuroscience research. Genetically encoded voltage indicators are a promising tool for this purpose, and recent developments have achieved high signal to noise ratio *in vivo* with 1-photon fluorescence imaging. However, these recordings exhibit several sources of noise that present analysis challenges, namely light scattering, out-of-focus sources, motion, and blood flow. We present a novel signal extraction methodology, Spike-Guided Penalized Matrix Decomposition-Nonnegative Matrix Factorization (SGPMD-NMF), which resolves supra- and sub-threshold voltages with high fidelity, even in the presence of correlated noise. The method incorporates biophysical constraints (shared soma profiles for spiking and subthreshold dynamics) and optical constraints (smoother spatial profiles from defocused vs. in-focus sources) to cleave signal from background. We validated the pipeline using simulated and composite datasets with realistic noise properties. We demonstrate applications to mouse hippocampus expressing paQuasAr3-s or SomArchon, mouse cortex expressing SomArchon or Voltron, and zebrafish spine expressing zArchon1.

## Introduction

A technology to measure the membrane potential of multiple neurons simultaneously in behaving animals would be a powerful tool for neuroscience research. Whereas calcium imaging (Farhi et al., 2019; Sofroniew et al., 2016) and extracellular arrays (e.g. neuropixels (Jun et al., 2017)) can report the spiking outputs of hundreds or even thousands of neurons, the sub-threshold dynamics are invisible to these methods. Sub-threshold voltages reflect synaptic and neuromodulatory inputs, as well as the activity of many endogenous ion channels. Precise spike waveforms are also useful to distinguish different spike types, e.g. complex vs. simple spikes, and subcellular spike modulations (Panzera and Hoppa, 2019). Accurate information on membrane voltages is essential for constraining detailed biophysical models of circuit function.

Several efforts in the last year introduced new genetically encoded voltage indicators (GEVIs): QuasAr3 (Adam et al., 2019), Archon (Piatkevich et al., 2019), Voltron (Abdelfattah et al., 2019), and ASAP3 (Villette et al., 2019). In combination with advanced optical instrumentation, these GEVIs achieved fluorescence voltage imaging from identified neurons *in vivo* (Bando et al., 2019). Pairing of optogenetic stimulation with voltage imaging can reveal the relative contributions of synaptic excitation and inhibition to the membrane voltage (Fan et al., 2020). These recordings, however, have substantial statistical and systematic noise sources that present a challenge for extraction of true neuronal voltage dynamics.

Voltage imaging *in vivo* presents far more stringent technical challenges than does calcium imaging. First, accurate detection of action potentials requires frame rates of ∼1,000 frames/s (or even higher for fast-spiking interneurons), vs. typically ∼10 frames/s or slower for Ca^2+^ imaging. Photons from voltage reporters are thus divided into ∼100-fold thinner time bins and consequently have ∼10-fold higher shot noise relative to signal, all else being equal.

Second, GEVIs typically yield 10-40% ΔF/F per spike, whereas the best Ca^2+^ indicators have several-fold increases in brightness for a single action potential, but are insensitive to subthreshold potentials (Chen et al., 2013; Tian et al., 2009). For millivolt-level subthreshold events GEVIs typically produce ΔF/F < 0.5%. Thus the subthreshold signals in voltage imaging can be as much as 100-fold smaller than the spike signals in Ca^2+^ imaging.

Third, the most advanced voltage imaging schemes rely on 1-photon optics (though there has been recent progress in 2P voltage imaging (Villette et al., 2019)). Consequently, 1P voltage imaging experiments are susceptible to optical crosstalk between distinct in-focus signal sources; and also from out-of-focus cells, blood flow, or background autofluorescence.

Fourth, subthreshold voltages often have strong correlations between cells, while spiking tends to be less correlated (Lampl et al., 1999). Thus the true voltage signal of interest is often correlated with sources of crosstalk.

Fifth, voltage signals only originate at the cell membrane, a nanometers-thick 2D manifold, whereas Ca^2+^ signals come predominantly from the cytoplasm throughout the cell body. Voltage measurements are thus far more sensitive to motion artifacts or to misalignment of illumination, sample, and detection. During behavioral tasks, motion artifacts may also correlate with true signal.

Due to the confluence of these factors, signal extraction for voltage imaging presents unique challenges. Correlated signal and noise dynamics confound well established signal extraction techniques, such as principal components analysis-independent components analysis (PCA-ICA) (Mukamel et al., 2009), that assume statistical independence between distinct sources. Advanced image demixing techniques for *in vivo* 2P Ca^2+^ imaging data are primarily focused on identifying spiking events, and thus can safely neglect many sources of noise (Pnevmatikakis et al., 2016).

Recently, a joint penalized matrix decomposition (PMD) and non-negative matrix factorization (NMF) approach has been proposed to denoise and demix voltage imaging data (Buchanan et al., 2019). This method can extract cell signals that have high signal to noise ratio (SNR) from *in vitro* voltage imaging movies, where motion artifacts, blood flow, light scattering, and temporally-varying background can all be ignored.

Here, we build upon the PMD-NMF pipeline to account for fluctuating background dynamics that might be correlated with the desired in-focus neural signals. We call the augmented pipeline Spike-Guided PMD-NMF (SGPMD-NMF). Our procedure takes advantage of several robust statistical structures within voltage imaging data. First, action potentials have lower correlations between cells and to noise sources than do slower subthreshold signals. We thus extract the spatial footprints of cells from spikes alone. We then apply the same spatial footprints to extract the slower subthreshold voltage dynamics. Spiking and subthreshold footprints coincide when cells are electronically compact, a reasonable assumption for studies performed with soma-localized GEVIs. The tradeoff is that the algorithm requires cells to spike. Cells that show purely subthreshold dynamics during the analyzed epoch are not detected.

Second, we use the fact that the spatial profiles of out-of-focus background sources are more dispersed and smooth compared to the spatial profiles of in-focus cell signals. By optimizing the smoothness of the background we determine how to apportion low-frequency fluorescence signals among signal and background, even when these two sources overlap in space and are correlated in time.

Using these constraints, we accurately resolve the subthreshold dynamics of neurons recorded *in vivo*. We validate our methods with simulated data containing realistic noise and where ground truth is known by construction. We further validate the method with composite movies created by summing real *in vivo* single-cell recordings. In these cases, we compare SGPMF-NMF with PCA-ICA and find that SGPMD-NMF produces traces that more closely resemble ground truth extracted from the individual component movies. Finally, we apply our pipeline to datasets from different organisms (mouse, zebrafish) and using different indicators (paQuasAr3-s, SomArchon, zArchon1, Voltron) to show the broad applicability of SGPMD-NMF. Code is available here: https://github.com/adamcohenlab/invivo-imaging. Analysis was conducted using both Matlab and Python.

## Model and Methods

The SGPMD-NMF algorithm comprises two steps: 1) Denoising, and 2) Demixing. The sub-steps are shown in Figure 1.

**Figure 1.**
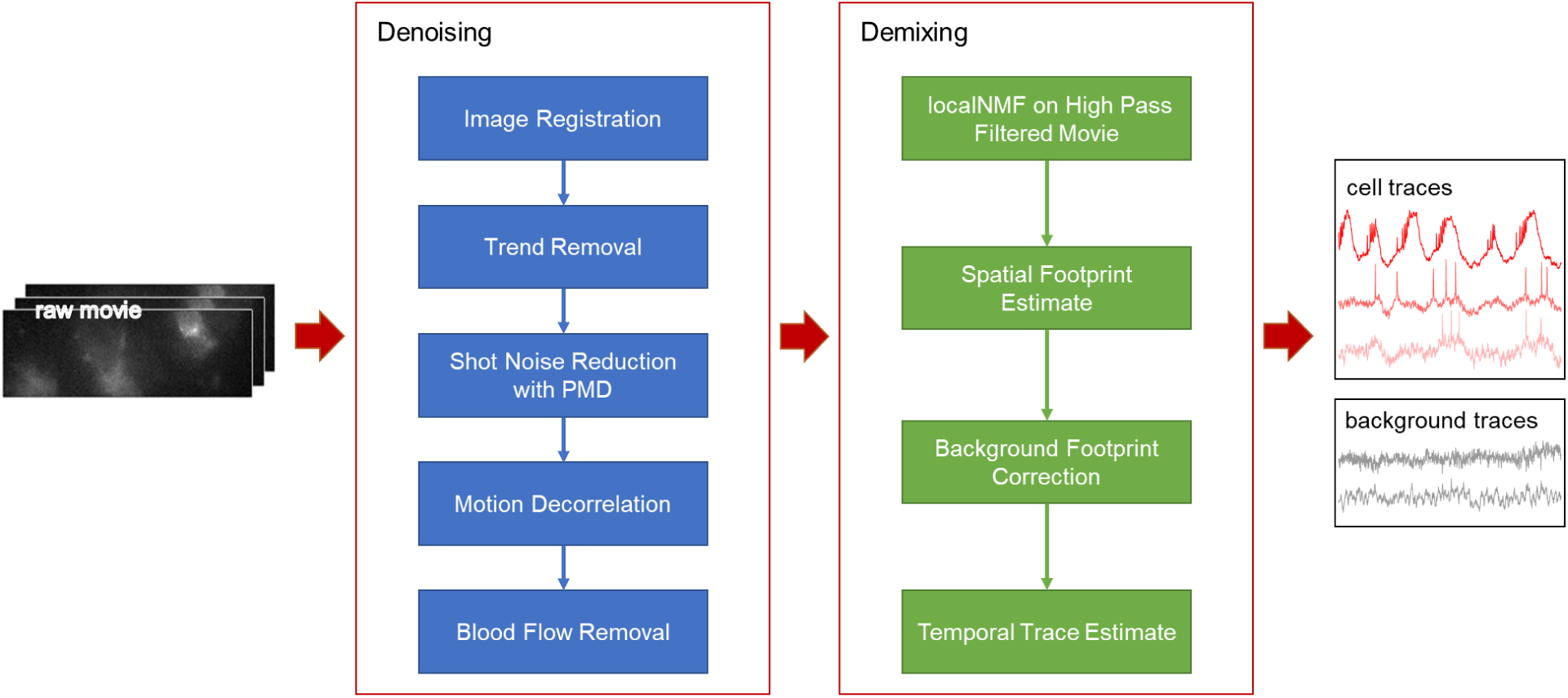
Pipeline for denoising and demixing *in vivo* voltage-imaging data. The Denoising steps (blue) comprise a set of distinct corrections for the sources of statistical noise and systematic artifacts that can arise *in vivo*. The Demixing steps (green) use action potentials to identify the cell footprints, and the differing spatial profiles of in-focus vs. out-of-focus sources to apportion subthreshold dynamics between cells and background.

### Denoising

The denoising steps address several distinct sources of noise: motion artifacts, photobleaching, shot noise, and blood autofluoresence. The raw dataset is first corrected for in-plane motion using NoRMCorre (Pnevmatikakis and Giovannucci, 2017). This step aligns the neuron locations between frames of the movie. The function also returns the horizontal (x) and vertical (y) displacements of each frame relative to a reference frame.

Photobleaching often does not follow simple monoexponential kinetics, particularly when the illumination is nonuniform such that different molecules experience different local illumination intensities. Rather than using an exponential fit, each pixel in the registered movie is corrected for photobleaching with a 3rd-order spline-based detrending fit. In selecting the detrending interval one must select an interval short enough to correct for the fastest photobleaching transients, but not so short that sub-threshold depolarizations (e.g. network up states) are removed. Typically, 5 s is appropriate.

Shot noise is removed from the movie via a spatially-localized PMD approach (Buchanan et al., 2019). This approach uses the fact that shot noise is uncorrelated between pixels, whereas true voltage signals typically extend over multiple contiguous pixels. The PMD approach calculates local filters that preserve the correlational structure in the movie. To achieve reasonable run time for long movies, we typically apply the PMD algorithm to a subset of contiguous frames selected from the full movie. PMD returns spatial filters, which are then applied to the full movie to obtain the denoised version of the full movie. This approach allows the PMD algorithm to run at a fixed time even as the movie duration increases.

NoRMCorre did not completely correct for motion artifacts for two reasons. First, it did not correct for small out-of-plane z motions which affected the focus of the image. Second, it did not correct for relative motion of sample and spatially structured fluorescence excitation light. Structured illumination can dramatically improve SNR in stationary samples by minimizing out-of-focus fluorescence (Adam et al., 2019; Chien et al., 2017; Fan et al., 2020). However, brain motion shifts the signal sources relative to the illumination, leading to modulation of fluorescence which is not corrected by rigid-body image translations. To reduce the effects of these artifacts, we employed a generalized linear model to project motion-correlated signals out of each pixel in the denoised movie. We removed from each pixel any components of the signal correlated with the regressors [x(t), y(t), x^2^(t), y^2^(t), x(t)y(t)], where x(t) and y(t) represent the motion traces calculated by NoRMCorre. Finally, to reduce contamination from blood flow, the software pipeline includes functionality to manually select and mask regions containing blood vessels.

### Demixing

After cleaning the noise from the movie, we approached the “demixing” problem of separating individual neurons and background spatial components, or footprints. We took advantage of the greater statistical independence of spiking compared to subthreshold voltages to identify the single-cell footprints. To reduce analysis time for large FOVs, we applied a 2×2 binning to the denoised movie before demixing. First, we applied a sliding window temporal high-pass filter (cutoff 10 ms) to each pixel in the denoised movie to remove the low-frequency subthreshold signals. The PMD denoising step enables us to isolate spiking signals at the single-pixel level with a temporal high-pass filter. Without PMD denoising, single-trial single-pixel spike finding would be impeded by temporally-uncorrelated shot noise.

The high-pass filtered movie was then processed by a local non-negative matrix factorization (localNMF) demixer, which detected cell profiles by identifying sets of highly-correlated, spatially contiguous pixels (Buchanan et al., 2019). The initial demixer output, ***A***_**0**_, consisted of the *d* × *n* matrix of cell footprints, where *d* is the number of pixels in a frame and *n* is the number of spiking cells. Footprints from nearby cells were allowed to overlap. A threshold was applied to the initial estimates of the cell footprints, ***A***_**0**_, to set small values to zero and thereby to delimit the spatial support of the cells.

We then sought to identify the time-course of the background dynamics. PCA was applied to the original movie (after denoising, before high-pass filtering), with analysis restricted to pixels not in the thresholded ***A***_**0**_. A user-specified number of background components was selected, typically ≤ 10. These components represent contributions to the fluorescence from out-of-focus cells, breathing artifacts, or residual motion artifacts. We assumed that the time-course of the background contributions to each cell would be a linear combination of the off-cell background dynamics.

The next step was to apportion the on-cell dynamics between background and signal. Initially the background footprints were set to zero under all the cells. We then used a fast hierarchical alternating least squares (fastHALS) algorithm (Friedrich et al., 2017) to fit background weights and cell temporal components to the full denoised movie (including the low-frequency dynamics). By using cell spatial weights initialized from the spiking-only data, we ensured that the extracted low-frequency dynamics had the same spatial footprints as the spiking, a consequence of the electrotonic compactness of cell bodies. After iterating the least squares fit, the output gave updated cell spatial footprints, ***A***_**1**_, and a fit for the *d* × *r* matrix of background spatial footprints, ***B***_**0**_, where *r* is the estimated rank of the background.

At this step, ***B***_**0**_ typically contained contamination from ***A***_**1**_, i.e. the spatial maps of the background had structure that resembled the in-focus cells. This crosstalk implied that subthreshold signals were not optimally apportioned between in-focus cells and background. To remedy this crosstalk, we sought to remove the spatial footprints of the cells from the images of the background. Here we used the fact that out-of-focus background was spatially smoother than in-focus signal.

We created updated background spatial components ***B***_**1**_ = ***B***_**0**_− ***A***_**1**_**× *W***_***opt***_, where ***W***_***opt***_ is a *n* × *r* matrix of weights selected to maximize a measure the spatial smoothness in each column of ***B***_**1**_.

We chose each column, *j*, of the weight matrix by optimizing the following through gradient descent:

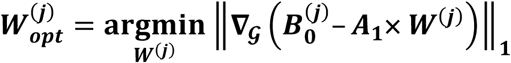

where ∇_𝒢_ is the nearest-neighbor discrete approximation to the gradient operator and

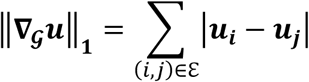

where pixels (*i, j*) are in the edge set ε when the pixels are adjacent. The gradients at points where the objective was non-differentiable were set to zero for the optimization procedure.

With the updated footprints, we implemented a least-squares estimate of membrane voltage by regressing the cell spatial components and updated background images on the full denoised and motion-corrected (but not high-pass-filtered) movie. This process gave the final output traces, which isolated cells and accurately separated their subthreshold signals from the background dynamics. A key feature of this algorithm is that it did not impose statistical independence of cell and background temporal sequences or spatial profiles.

Both the PMD denoising and localNMF demixing steps scale linearly with number of movie frames, so to reduce the time for analysis, in our implementation, we run the PMD denoising and localNMF demixing steps on only a subset of the frames of the movie (typically 6000 frames for denoising and 10,000 frames for demixing) to obtain spatial filters (for denoising) or cell and background spatial footprints (for demixing), which are then applied to the whole movie. This step reduces the total time of analysis for a 96 × 284 pixel movie of 60,000 frames to under 20 minutes on the Harvard FASRC Cannon cluster.

## Results

### SGPMD-NMF pipeline reliably recovers cell signals in the presence of correlated background

We verified the pipeline on simulated data containing two partially overlapping disk-shaped cells and a spatially heterogeneous time-varying rank-1 background (Fig. 2A). To impose correlated sub-threshold dynamics, we constructed sets of three sub-threshold waveforms with a specified 3×3 cross-correlation matrix, and then assigned one waveform to each of the cells and one to the background. We then added spikes (2 ms wide) atop the waveforms at independent Poisson-distributed intervals. Spike height and subthreshold power spectrum were selected to approximately correspond to *in vivo* recordings. Poisson-distributed shot noise was added to each pixel, and the SNR was adjusted by varying the overall brightness.

**Figure 2.**
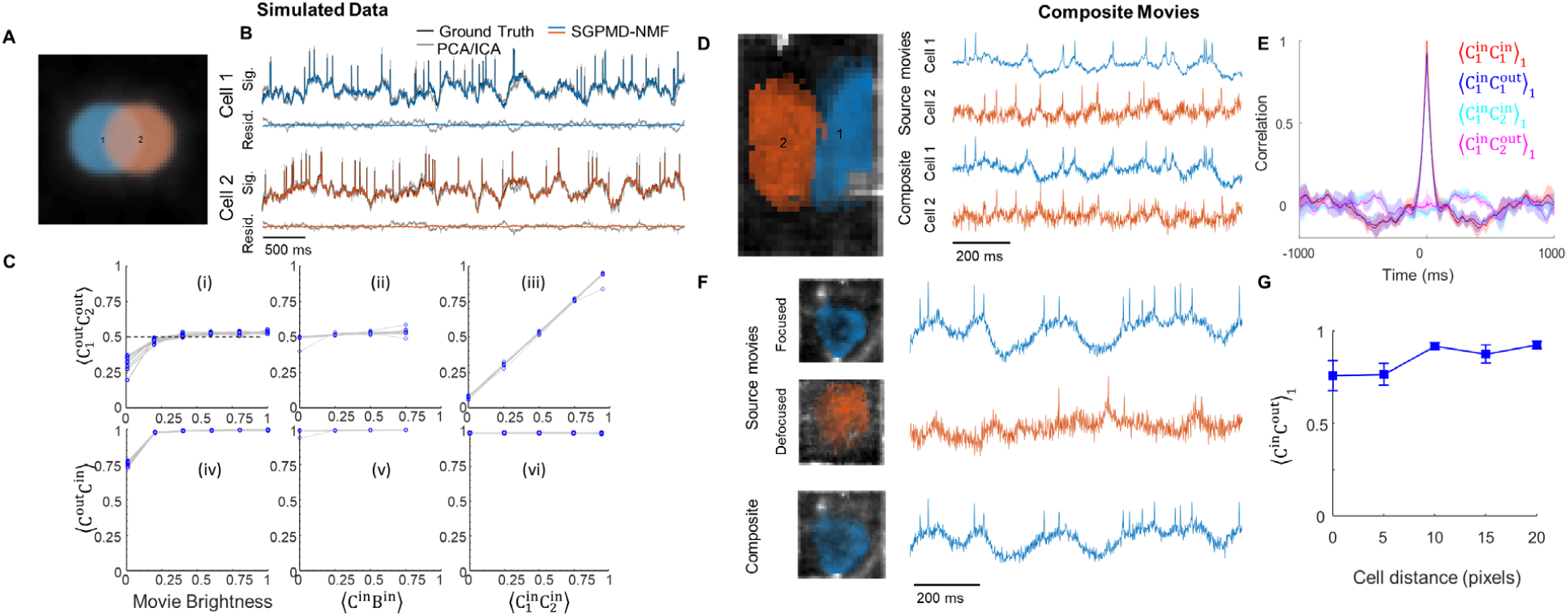
Validation of the SGPMD-NMF algorithm on simulated data (left) and composite movies (right). A) Example field of view comprising two simulated cells with spatial overlap and correlated subthreshold dynamics, and a broad background whose dynamics were also correlated with each of the cells. B) Comparison of signal extraction via PCA-ICA vs. SGPMD-NMF. The signals extracted via SGPMD-NMF were substantially closer to the ground truth than were the signals extracted via PCA-ICA. C) Quantification of SGPMD-NMF performance as a function of signal characteristics. Here ⟨*XY*⟩ is the cross-correlation of *X* and *Y. C*_1_ and *C*_2_ are the signals of the two cells, *B* is the background. When subscripts are omitted, the calculation is averaged over both cells. In panels i, ii, iv, v, 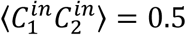. In panels i, iii, iv, vi, ⟨*C*^*in*^*B*^*in*^⟩ = 0.5. In panels ii, iii, v, vi, movie brightness = 1. D) Left: Composite movies were formed by adding two separately acquired single-cell movies (mouse hippocampus expressing paQuasAr3-s) with a 20 pixel lateral offset between the cells. Right: signals were extracted from the source movies individually and from the composite movie jointly. E) Temporal cross-correlations of input and output traces showed good fidelity of extracted relative to input traces. Here ⟨*XY*⟩_1_ is the cross-correlation of *X* and *Y* normalized by its value at lag 1 time-step (see Statistical Methods for details). F) Left: A second set of composite movies were formed by adding two separately acquired single-cell movies of the same cell (mouse hippocampus expressing paQuasAr3-s), one with the cell in-focus and one with the cell 20 μm out of focus. The lateral offset between the two cells ranged from 0 pixels to 20 pixels. Right: signals were extracted from the source movies individually and from the composite movie jointly. In the composite movie, only the in-focus cell was extracted as a cell signal, and the out-of-focus cell was treated as background. G) Correlation of output traces with corresponding input traces as a function of overlap between the two cells. Error bars show mean ± s.e.m.

We explored the ability of the algorithm to retrieve the input signals as a function of the SNR and the sub-threshold correlations. First, we compared the performance of PCA-ICA and SGPMD-NMF (Fig. 2B). In the displayed example, we analyzed a simulation where the subthreshold correlations between cells and between each cell and the rank-1 background were all 0.5. When all traces were normalized (mean of 0 and standard deviation of 1), the root mean square error of the traces extracted by PCA-ICA was 0.44, while for SGPMD-NMF it was 0.11. Furthermore, the correlation between the extracted PCA-ICA traces for the two cells was 0.19, while the correlation between the extracted SGPMD-NMF traces was 0.53, close to the ground-truth value of 0.5. This lower correlation from PCA-ICA compared to ground truth is expected, as ICA explicitly searches for independent components. Together these results demonstrate that SGPMD-NMF offers superior performance compared to PCA-ICA in extracting the subthreshold dynamics.

We then tested the ability of the SGPMD-NMF algorithm to preserve the correlations between Cell 1 and Cell 2, and between each cell and its respective ground truth, under different conditions of SNR and subthreshold correlations (Fig. 2C). Under all but the most stringent conditions the algorithm extracted the input signals with high fidelity. A movie brightness of 1 represented the SNR of a typical *in vivo* recording from (Adam et al., 2019).

### SGPMD-NMF pipeline preserves correlation structure in overlapping cell movies

A challenge with validating the algorithm on real-world data is that it is technically infeasible to make whole-cell patch clamp recordings from multiple neurons *in vivo* while simultaneously performing voltage imaging. Thus, there is no ground truth data against which to compare the outputs of SGPMD-NMF.

To address this challenge, we created composite movies by adding together real *in vivo* voltage imaging movies of well-isolated hippocampal neurons expressing paQuasAr3-s. By adjusting the lateral offset of the two cell images we could control the degree of spatial overlap in the composite movie. This approach tested the algorithm beyond the “worst-case” of *in vivo* crosstalk because two real cells could never occupy the same volume.

We extracted the single-cell voltage traces from the individual recordings (giving “input” traces), and then applied SGPMD-NMF to the composite recordings (giving “output” traces, Fig. 2D). We performed the calculation for four pairs of cells, giving four trials. The correlation between the input traces was near zero, as expected for recordings taken at different times and different fields of view. In *in vivo* recordings, the distance of closest approach of in-focus cells is approximately 1 cell diameter, corresponding to 20 pixels in our movies. At this separation, the lag-1 cross-correlation between the output traces of the two cells was ⟨*C*_1_^*out*^*C*_2_^*out*^⟩_1_ = 0.17 ± 0.17 (mean ± s.e.m.), not significantly different from zero (see Statistical Methods for rationale for using lag-1 correlations). The mean lag-1 correlation between each output trace and its corresponding input trace at this separation was ⟨*C*^*in*^*C*^*out*^⟩_1_ = 0.92 ± 0.02 (mean ± s.e.m.). The temporal auto- and cross-correlation functions of the output traces closely matched the corresponding functions of the input traces (Fig. 2E).

We next tested algorithm performance in the presence of out-of-focus spatially overlapping sources. We added *in vivo* voltage imaging movies of well-isolated hippocampal neurons expressing paQuasAr3-s to subsequent recordings of the same cells taken with 20 μm defocus (Fig. 2F). Addition of an out-of-focus background cell with a lateral offset of 20 pixels did not substantially perturb the extracted waveform of the in-focus cell (⟨*C*^*in*^*C*^*out*^⟩_1_ = 0.93 ± 0.02, mean ± s.e.m., *n* = 4 pairs, Fig. 2G). Shifting the out-of-focus cell to directly beneath the in-focus cell lowered this correlation to ⟨*C*^*in*^*C*^*out*^⟩_1_ = 0.76 ± 0.08.

### SGPMD-NMF pipeline works for multiple species and reporters

After validating SGPMD-NMF in simulations and composite datasets, we then applied the pipeline to analyze multi-cell *in vivo* recordings acquired with different species, brain regions, cell types, reporters, and imaging modalities (Table 1, Methods). The reporters and imaging systems have all been published previously (Abdelfattah et al., 2019; Adam et al., 2019; Fan et al., 2020; Piatkevich et al., 2018). The zebrafish data were acquired on a previously unpublished transgenic line expressing zArchon1 under control of the Vglut2a enhancer (Experimental Methods).

**Table 1.** *In vivo* datasets analyzed via SGPMD-NMF. Diverse data-sets included different species (mouse, zebrafish), regions of the central nervous system (hippocampus, cortex, spinal cord), reporters (paQuasAr3, SomArchon, zArchon1, Voltron), and imaging modalities (structured illumination, holographic, wide-field epifluorescence, and light sheet).

| Brain region | Cell type | Reporter | Imaging modality | Reference |
| --- | --- | --- | --- | --- |
| Mouse hippocampus | Oriens interneurons and pyramidal cells | paQuasAr3 (Fig. 2)<br>SomArchon (Fig. 3) | Micromirror-based soma-targeted structured illumination | Experimental setup: (Adam et al., 2019)<br>Reporter: (Piatkevich et al., 2019) |
| Mouse cortex L1 | Interneurons (Ndnf<sup>+</sup>) | Voltron | Wide-field epifluorescence | (Abdelfattah et al., 2019) |
| Mouse cortex L1 | Interneurons (5-HT<sub>3A</sub>R<sup>+</sup>) | SomArchon | Holographic peripheral membrane-targeted structured illumination | (Fan et al., 2020) |
| Zebrafish spinal cord | Excitatory (Vglut2a<sup>+</sup>) | zArchon1 | Light sheet | Transgenics: Unpublished<br>Reporter: (Piatkevich et al., 2018) |

Videos 1–4 show each step of the SGPMD-NMF pipeline for each of the preparations. SGPMD-NMF identified spiking cells and separately extracted time-dependent signals from in-focus cells (Figs. 3–6) and from multiple background components (Figs. S1–S4). The videos show that the initial denoising steps of SGPMD-NMF substantially reduce speckle, motion, and blood flow artifacts. The movies of the residuals (denoised movie minus signal and background) show that most sources of variation under spiking cells are accounted for in either signal or background; and that the extracted waveforms of the spiking cells in the signal look biologically plausible (e.g. elevated spike rates during periods of subthreshold depolarization). The background movie is spatially smooth near the identified cells and shows low-frequency temporal dynamics that are highly correlated across many pixels.

**Figure 3.**
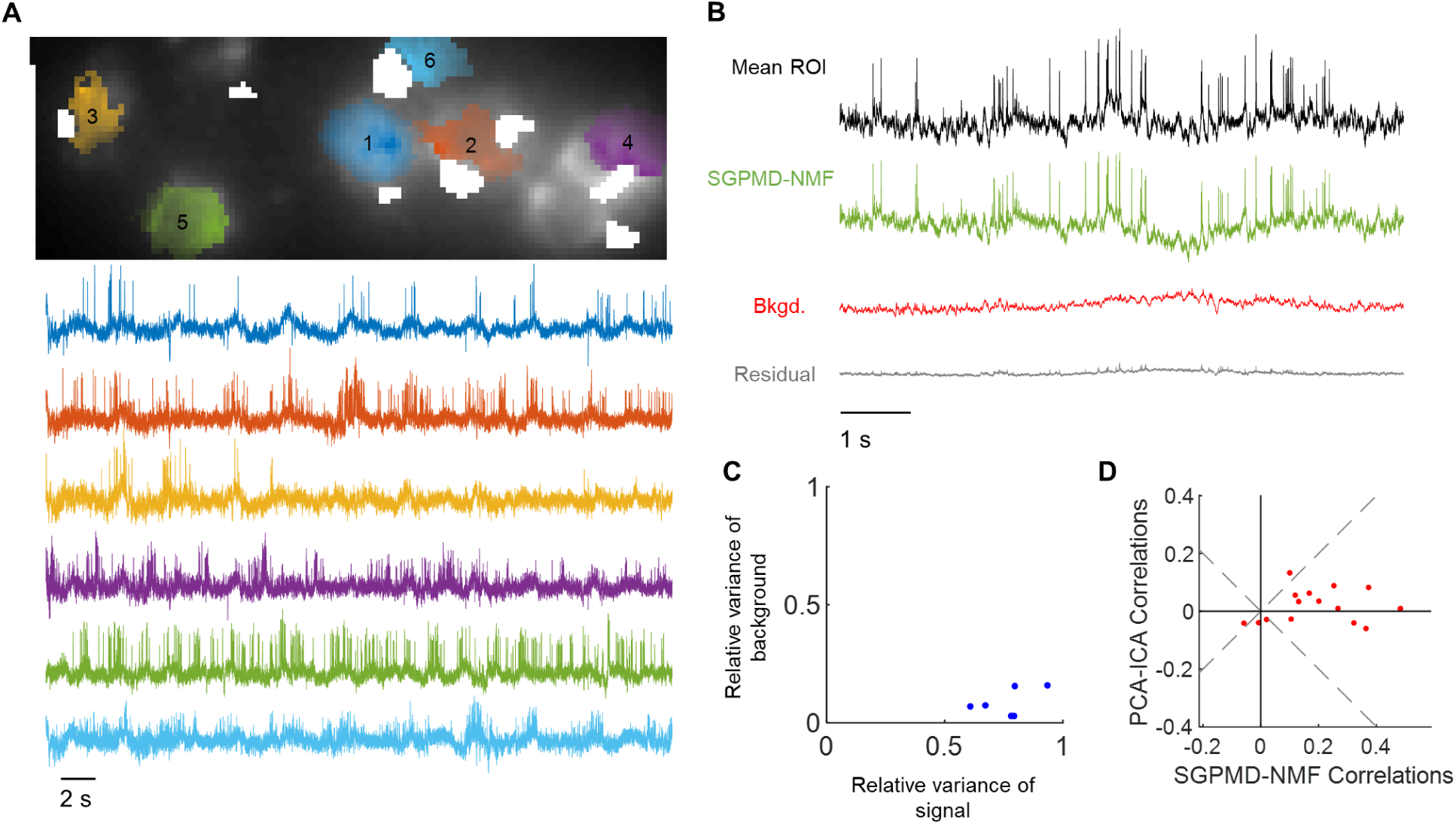
Voltage imaging in mouse hippocampus CA1 using SomArchon. The cells expressed SomArchon and were imaged via micromirror-based soma-targeted structured illumination. A) Top: Image of the field of view, showing dense neurons as occurs in the pyramidal cell layer of CA1. The cell footprints are overlaid. Regions contaminated by blood flow are masked in white. Bottom: extracted single-cell traces. Subthreshold depolarizations clearly coincided with elevated spike rates, giving confidence that the sub-threshold waveforms reflect membrane potential. B) Average across pixels in cell 5 in the denoised movie (Mean ROI), SGPMD-NMF reconstructed signal movie (SGPMD-NMF), reconstructed background movie (Bkgd.) and residual movie (Residual). C) Scatter plot of the relative variance of each cell background vs. signal. D) Comparison of the pairwise cell-cell cross-correlations between SGPMD-NMF and PCA-ICA. Most (12 of 15) pairwise correlations had a smaller magnitude for PCA-ICA vs. SGPMD-NMF.

**Figure 4.**
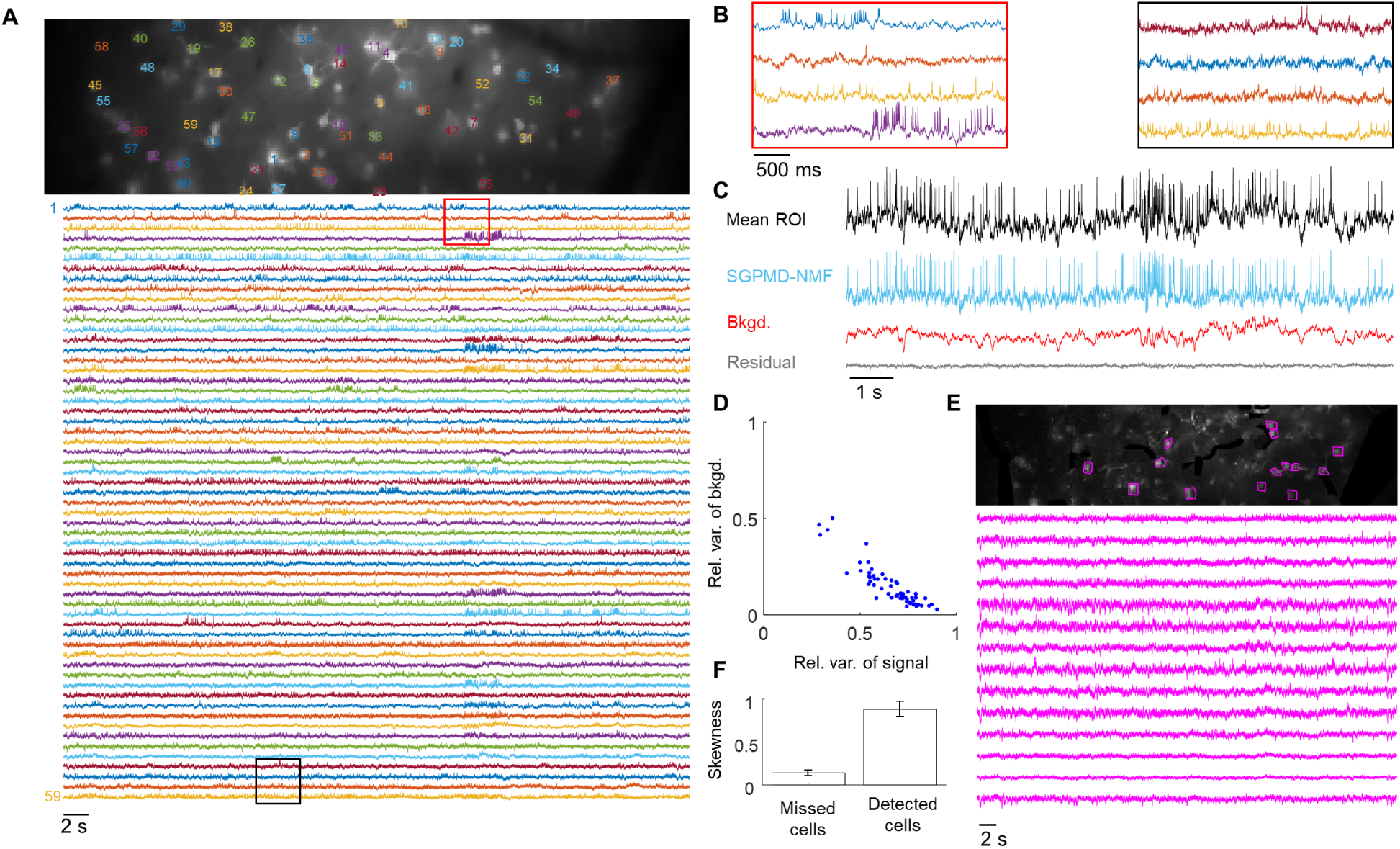
Voltage imaging in mouse cortex L1 using Voltron. The cells expressed Voltron and were imaged via wide-field epifluorescence. A) Top: Image of the field of view. The cells are labeled at the centroid of their footprints. Bottom: extracted single-cell traces. Insets shown in B) are marked in red and black. B) Inset of eight single cell traces over a window of approximately 3 s. C) Average over pixels in cell 6 in the denoised movie (Mean ROI), SGPMD-NMF reconstructed signal movie (SGPMD-NMF), reconstructed background movie (Bkgd.) and residual movie (Residual). D) Scatter plot of the relative variance of each cell background vs. signal. Anticorrelation between relative variance of background vs signal is due to the facts that together background and signal account for >99% of the total variance. If background and signal were uncorrelated, they would fall along the line *x* + *y* = 1. Deviations below the line *x* + *y* = 1 indicate positive correlation between signal and background. E) Top: Standard deviation image of the reconstructed sum of background and residual. Magenta masks indicate 14 manually selected bright spots, or “missed cells.” Bottom: Mean ROI (on denoised movie) of the 14 missed cells. These traces showed little or no spiking activity. F) Skewness of temporally high-pass filtered mean ROI traces (on denoised movie) of the 14 missed cells and 59 detected cells. Skewness provides a measure of spiking activity relative to baseline noise. Error bars show mean ± s.e.m. The missed cells displayed substantially less spiking activity compared to the detected cells.

**Figure 5.**
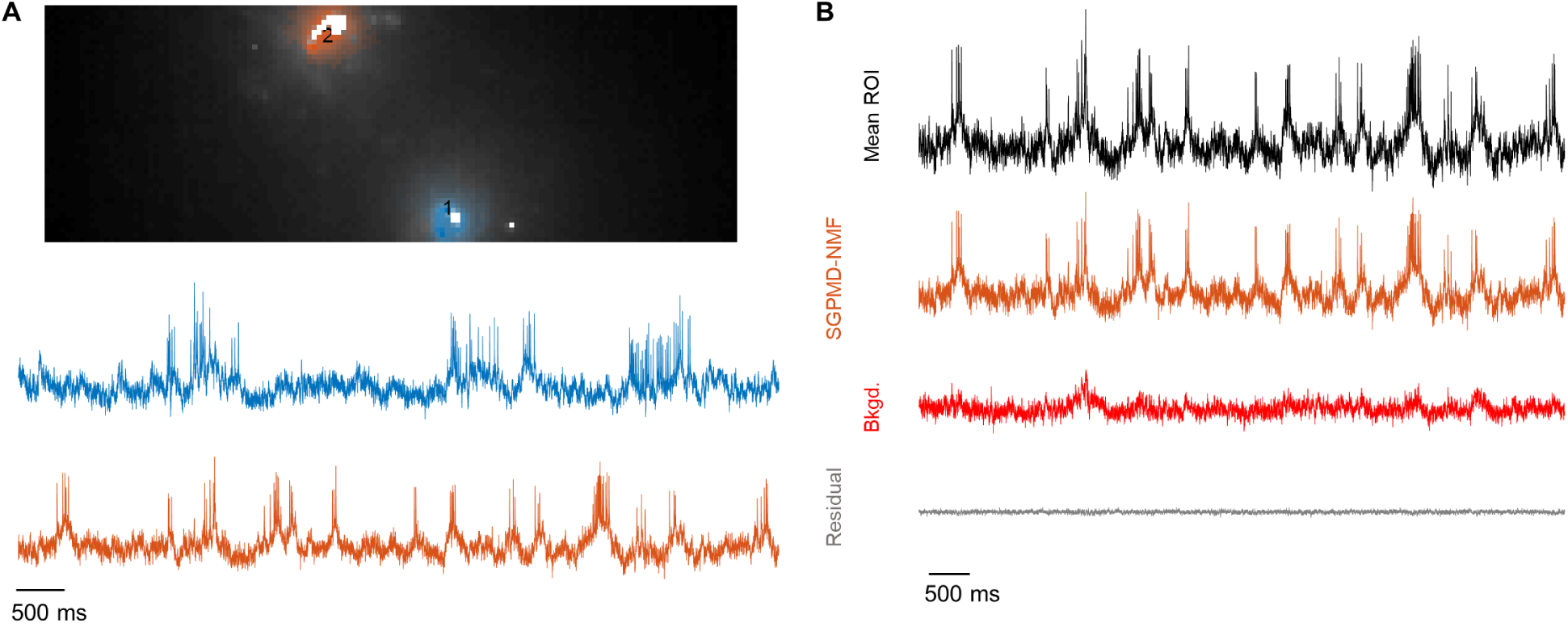
Voltage imaging in mouse cortical L1 using SomArchon. The cells expressed SomArchon and were imaged via holographic membrane-targeted illumination. A) Top: Image of the field of view, showing well-separated neurons as occurs in L1. The cell footprints are overlaid in blue and orange. Regions contaminated by blood flow are masked in white. Bottom: extracted single-cell traces. Elevated spike rates clearly reside atop subthreshold depolarizations, giving confidence that the sub-threshold voltages are at least approximately correct. B) Average across pixels in cell 2 in the denoised movie (Mean ROI), SGPMD-NMF signal (SGPMD-NMF), background (Bkgd.) and residual (Residual).

**Figure 6.**
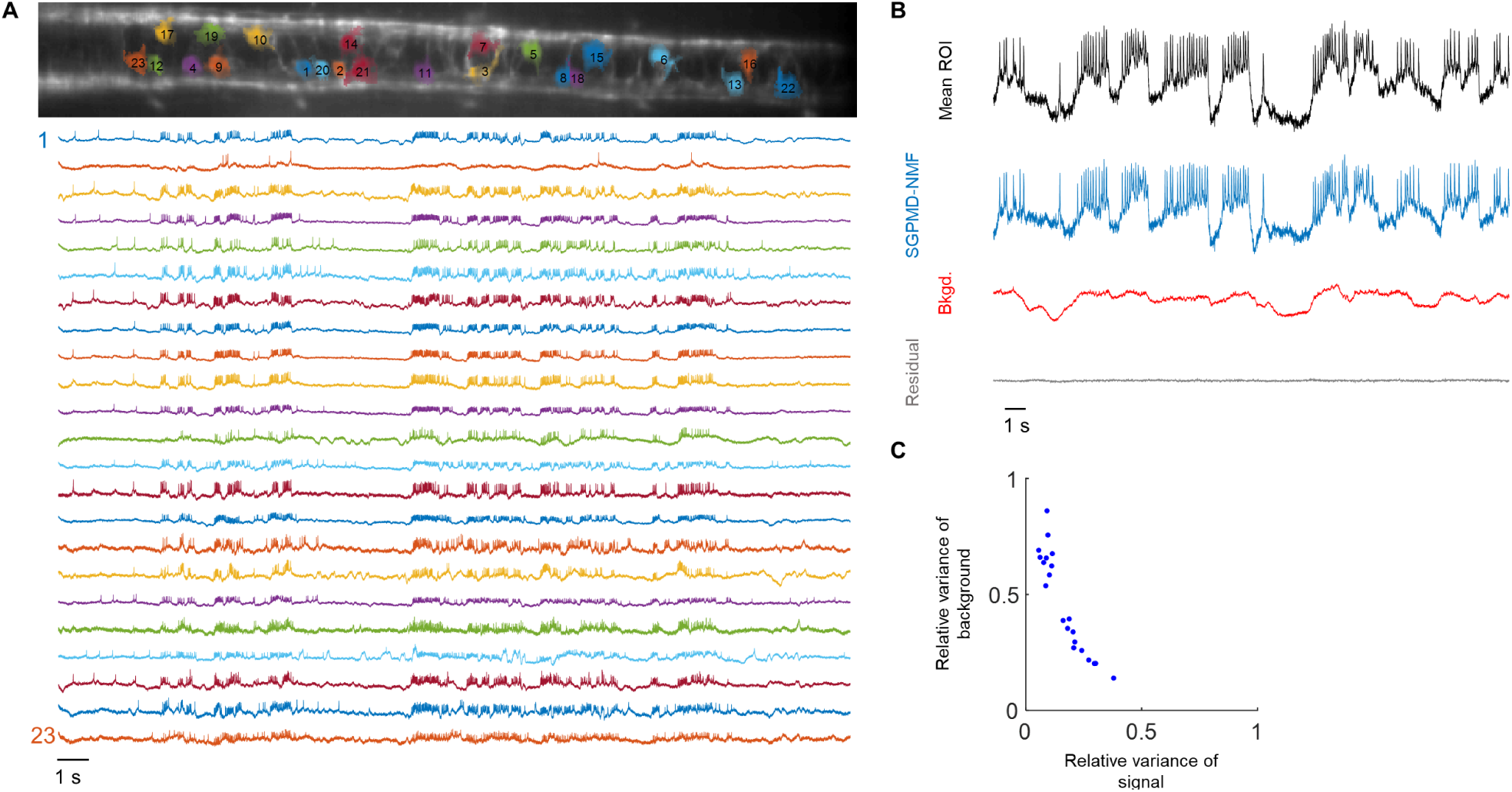
Voltage imaging in zebrafish spinal cord using zArchon1. The cells expressed zArchon1 and were imaged via light sheet fluorescence microscopy. A) Top: Image of the field of view. The cell footprints are overlaid. Bottom: extracted single-cell traces. B) Average over pixels in cell 8 in the denoised movie (Mean ROI), SGPMD-NMF signal (SGPMD-NMF), background (Bkgd.) and residual (Residual). C) Scatter plot of the relative variance of each cell background vs. signal. As in Fig. 4D, relative variance of background vs signal is anticorrelated.

To assess the performance of SGPMD-NMF, we compared the SGPMD-NMF outputs to what one would obtain from a flat average across each cell footprint after denoising, i.e. the mean of the ROI (Figs. 3B, 4C, 5B, 6B). The ROI-based signals contained all sources of background, while SGPMD-NMF traces isolated cell signals from background. To quantify the comparison, we quantified the ratios of the variances of the SGPMD-NMF signal, background, and residual traces to the variance of the mean ROI trace (Table 2). Most (> 99%) of the variance of the denoised movie within the cell ROIs was explained by a combination of signal and background, demonstrating that SGPMD-NMF accounted for almost all time-varying signals in the movies. Specifically, except for the zebrafish spinal cord dataset, the variance of the signal was on average larger than the variance of the background, indicating the cell regions had strong signal. For the zebrafish spinal cord dataset, we found that the background had a larger variance than the signal, due to strong contributions of blood flow to the background.

**Table 2.** Performance of SGPMD-NMF across different *in vivo* datasets. Relative variance was defined as the variance of the indicated SGPMD-NMF waveform divided by the variance of a flat average across the corresponding ROI. The relative variances do not sum to 1 because the distinct waveforms were not necessarily orthogonal.

| Brain region | Reporter | # of Cells | Relative Variance (mean ± s.e.m.) | | |
| --- | --- | --- | --- | --- | --- |
|  |  |  | Signal | Background | Residual |
| Mouse hippocampus | SomArchon | 6 | $0.76 \pm 0.05$ | $0.09 \pm 0.02$ | $0.01 \pm 0.00$ |
| Mouse cortex L1 | Voltron | 59 | $0.66 \pm 0.02$ | $0.15 \pm 0.01$ | $0.00 \pm 0.00$ |
| Mouse cortex L1 | SomArchon | 6 | $0.41 \pm 0.09$ | $0.35 \pm 0.07$ | $0.01 \pm 0.00$ |
| Zebrafish spinal cord | zArchon1 | 23 | $0.16 \pm 0.02$ | $0.52 \pm 0.06$ | $0.00 \pm 0.00$ |

Using the mouse hippocampus data, we also conducted a comparison of SGPMD-NMF to PCA-ICA (Fig. 3D). For the six cells identified in both methods, we calculated the 15 pairwise correlations between cell signals. For all but 3 cell pairs which happened to have low cross-correlations, the magnitude of correlation of the PCA-ICA signals was lower than that of the SGPMD-NMF signals. Based on the analysis with simulated data (Fig. 2), we infer that PCA-ICA likely underestimated cell-cell cross correlations in membrane voltage.

SGPMD-NMF analysis of the data from mouse cortex L1 using the reporter Voltron identified 59 spiking cells in a single FOV, including several overlapping footprints. However, the residual contained several objects that looked like bright cells but that were not picked up by the algorithm. To determine the reason for this, we studied the signals from ROIs around these cells. These cells all had very low SNR and lacked clearly discernable spikes. We quantified the number and amplitude of spikes via skewness. The mean skewness of the ROI signals of these cells after a 250 ms sliding window temporal high pass filter was 0.14 ± 0.03 (mean ± s.e.m.). For cells identified by SGPMD-NMF, the same metric was 0.88 ± 0.09. We inferred that SGPMD-NMF did not pick up these cells because they did not pass the initial spike finding step. To determine whether we could identify non-spiking cells, we analyzed a cropped portion of the dataset (Video 5). We employed a multi pass strategy for localNMF that identified additional cells with lower SNR (Buchanan et al., 2019).

**Video 1.**
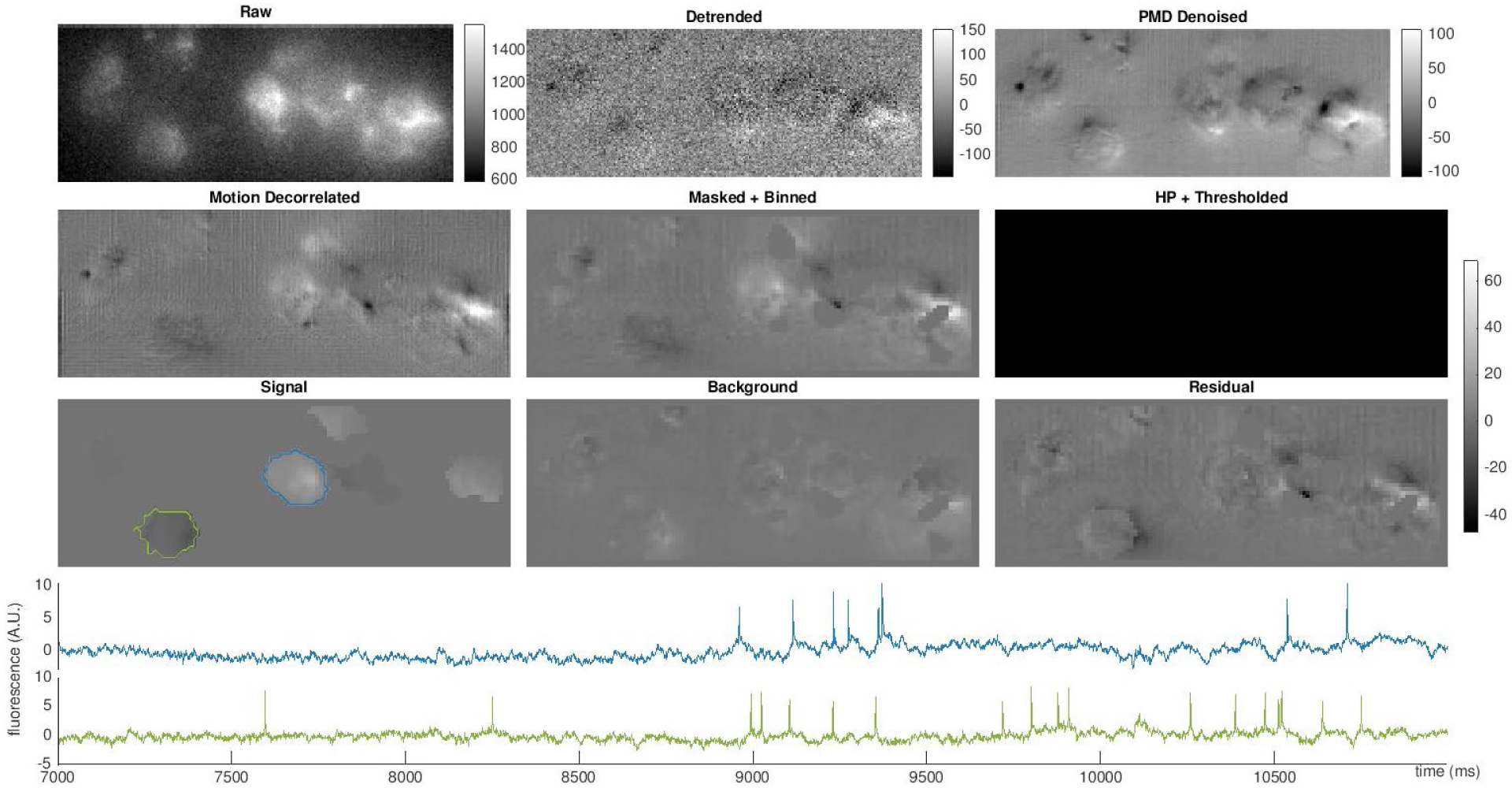
SGPMD-NMF applied to voltage imaging in mouse hippocampus CA1 using SomArchon. Top left: the recorded fluorescence data after NoRMCorre registration. Top middle: detrended data. Top right: PMD denoised data. Middle left: data with motion-correlated signals removed. Middle middle: data with pixels containing blood manually masked out. Middle right: temporally high-pass filtered and thresholded movie used for initialization of localNMF. Movie scale does not follow the color bar. Bottom left: SGPMD-NMF extracted signals reconstructed as a movie. Bottom center: SGPMD-NMF detected background reconstructed as a movie. Bottom right: residual movie (= masked data – signal – background). Bottom plots: time traces of two identified cells, marked in the signal movie. drive.google.com/open?id=114nBlwpTjVEpxft65KBlJB4qfoczZM_p

**Video 2.**
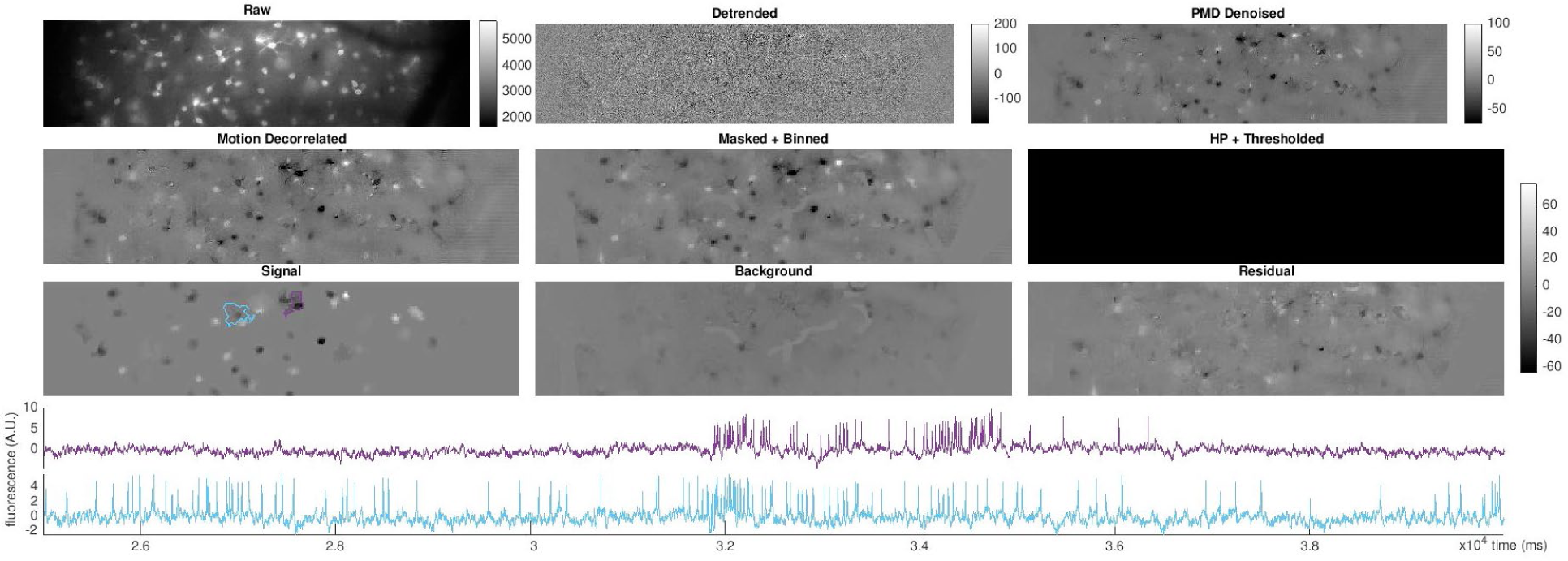
SGPMD-NMF applied to voltage imaging in mouse cortex L1 using Voltron. Panels are defined as in Video 1. The background and residual images contained cell-shaped objects. ROI analysis of these objects showed that they had minimal spiking activity during the recording, and hence were not detected by SGPMD-NMF. drive.google.com/open?id=14W_Xzj3VzhC6lnpC86WyHEfC9_o33Wzm

**Video 3.**
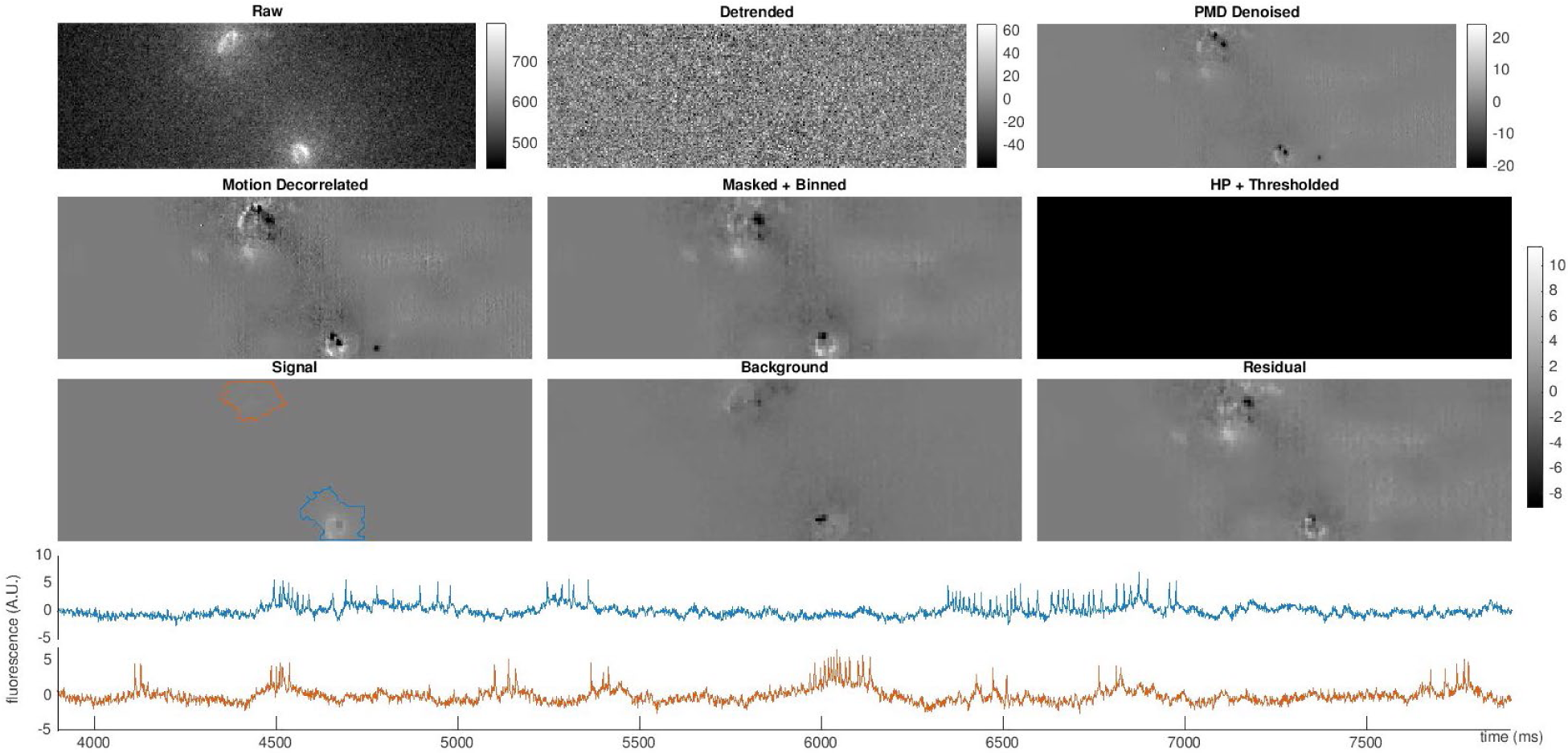
SGPMD-NMF applied to voltage imaging in mouse cortex L1 using SomArchon. Panels are defined as in Video 1. drive.google.com/open?id=14GRnAe7Szl5R2NMwhtfXkPgfK9HSFUgS

**Video 4.**
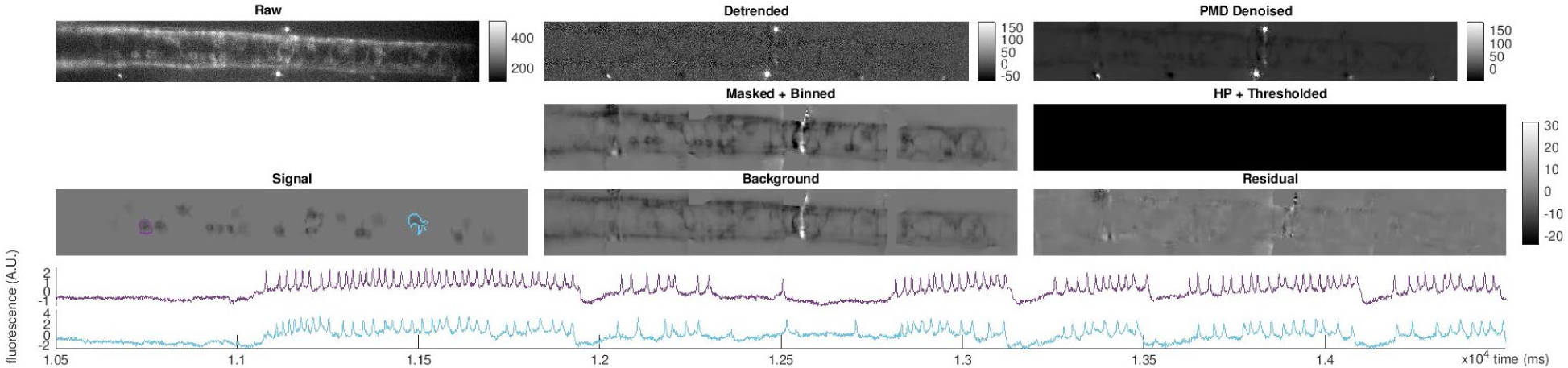
SGPMD-NMF applied to voltage imaging in zebrafish spinal cord using zArchon1. Same conventions as Video 1, but the motion correction step was not run for this dataset because the animal was paralyzed and there was minimal motion. drive.google.com/open?id=1qtCV1uyOmPl1Hjw3oGjt1xe5NtViQDKr

**Video 5.**
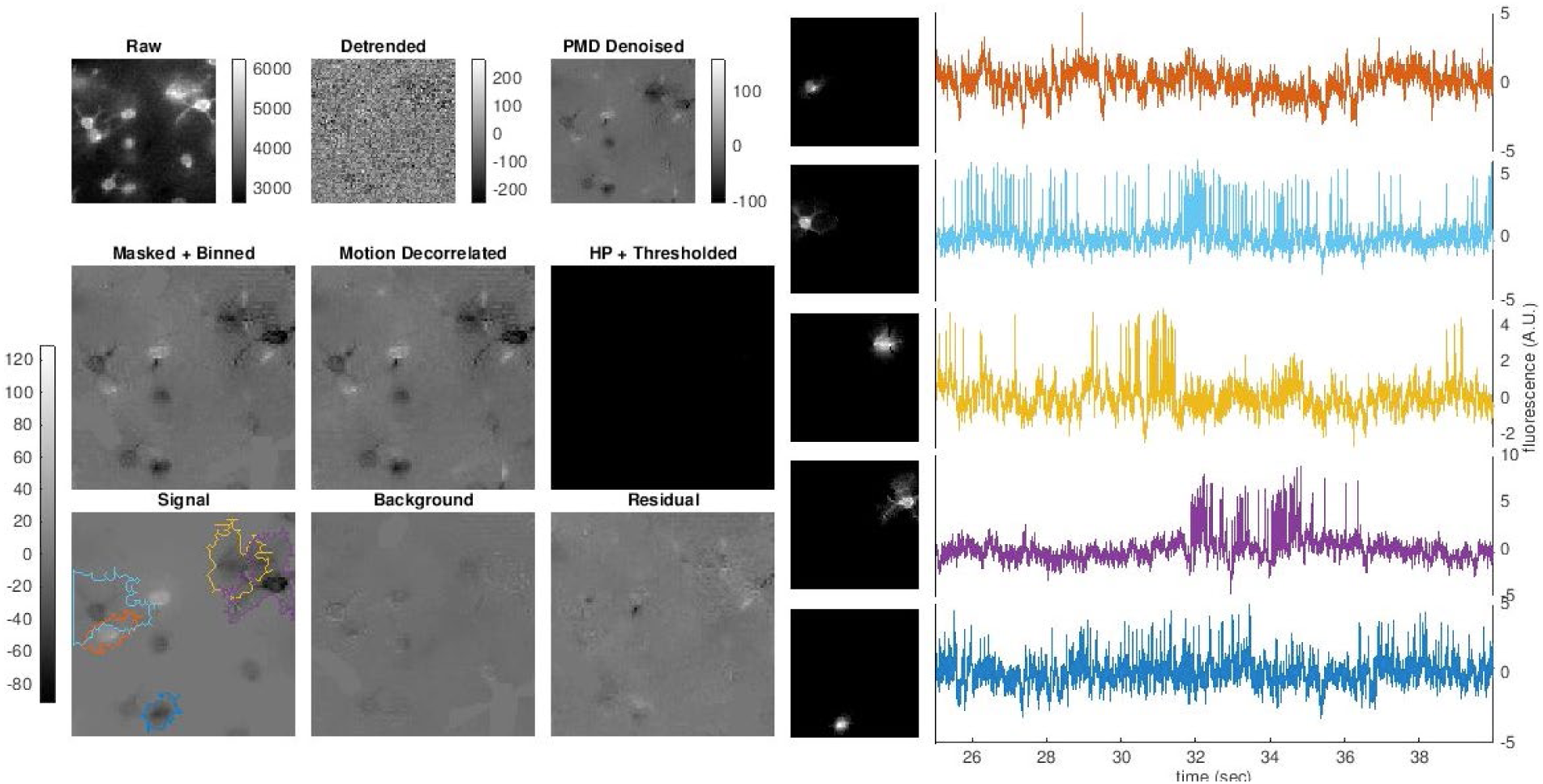
SGPMD-NMF applied to voltage imaging in mouse cortex L1 using Voltron (cropped FOV). Left: Same conventions for movies as Video 1. Right: Spatial footprints and temporal traces for five example cells in the FOV. Overlapping footprints are observed. drive.google.com/open?id=1r9Vg3smCverw1B4O5yP5pabQrrURb-HP

## Discussion

*In vivo* voltage imaging is a promising approach for studying the intercellular correlations in spiking and subthreshold dynamics. However, the data analysis must be performed with great care to avoid introducing spurious correlations or systematic artifacts. We have demonstrated a robust software pipeline for demixing voltage signals and background in complex, noisy tissues.

A further outstanding challenge is to identify strategies for creating masks that optimize the SNR of the extracted traces. Both shot noise and systematic noise (e.g. artifacts from blood flow, uncorrected motion, diffusing intracellular vesicles) vary over a cell. There are many combinations of pixel weights that give, on average, the correct voltage, but that differ in noise statistics. If the noise is independent across pixels, then the HALS approach optimizes the signal. When this assumption is not true, appropriate incorporation of a spatiotemporally correlated noise model could improve the quality of the extracted traces. Automated detection of blood vessels would also decrease manual labor in running the pipeline.

An independent voltage imaging spike detection pipeline, VolPy was recently introduced by (Cai et al., 2020). This pipeline is largely complementary to ours, as it focuses on extracting spike times, rather than accuracy of sub-threshold dynamics. VolPy uses a supervised neural network to find neurons based on summary images of the datasets. A similar approach could be adopted here to use cell morphology to identify both spiking and non-spiking cells as initial footprint guesses in the SGPMD-NMF demixing step; this will be an interesting direction to explore in future work. Compared to VolPy, SGPMD-NMF offers more effective demixing in several respects. First, denoising of temporally uncorrelated noise via PMD allows for improved accuracy in demixing (Buchanan et al., 2019). Second, SGPMD-NMF enables recovery of subthreshold signals and correlations with high fidelity by incorporating a background model that can have temporal dynamics correlated with cell signals. Without a proper method to account for the background, subthreshold signals of extracted cells will be inaccurate. Third, blood artifacts, which are not considered by VolPy, contaminate extracted cell signals as regions of blood flow often overlap with cell footprints. In many cases these blood artifacts are difficult to see in the raw data but become readily apparent after the denoising step used here.

With progress in 2P voltage imaging (Villette et al., 2019), challenges of optical crosstalk are expected to abate, but other challenges arise. Voltage signals only come from the intersection of the sub-micron 2P laser focus and the nanometers-thick cell membrane, leading to extreme sensitivity to motion artifacts. Furthermore, the stringent optical sectioning of 2P microscopy implies that all fluorescence must originate in a narrow equatorial belt of the cell, compared to 1P microscopy, which can average over the entire surface area of the cell. As a result, the requirements on per-molecule fluorescence are more demanding in 2P than in 1P microscopy, so photobleaching is more of a concern. Thus, revised algorithms may be needed for 2P voltage imaging. Our algorithm also assumed that cells were electronically compact. Applications to dendritic voltage imaging may require different approaches.

## Experimental Methods

### Mouse hippocampus expressing paQuasAr3-s data (used in composite movies)

See (Adam et al., 2019) for full details.

### Mouse hippocampus expressing SomArchon data

Optical and behavioral setup is the same as in (Adam et al., 2019). Reporter used is from (Piatkevich et al., 2019).

### Mouse cortex expressing SomArchon data

See (Fan et al., 2020) for full details.

### Voltron data

See (Abdelfattah et al., 2019) for full details.

### Zebrafish spine data

Recordings were done in 5 day-post-fertilization (dpf) transgenic zebrafish larvae expressing UAS:zArchon1-GFP under the control of vGlut2a:Gal4 (Satou et al., 2013). Larvae we paralyzed by immersion in 1 mg/ml α-bungarotoxin for ∼1 min and mounted in a drop of 1.5% low melting point agarose. Imaging was done on a custom light sheet microscope using a 639 nm red laser (MLL-FN-639, CNI lasers) to illuminate a 480 μm wide region with ∼300 mW of laser light. Images were collected through a low magnification high NA objective (XLPLN25XWMP2, Olympus) a 100 mm tube lens (TTL100-A, Thorlabs) and a 664 nm long pass filter. Images were recorded at 1kHz on a sCMOS camera (Hamamatsu Flash 4.0). During the recording, larvae were presented with a forward moving grating at 15 mm/s to induce swimming. To ensure naturalistic behavior, the ventral nerve root signal was electrophysiologically recorded at the same time and the fictive swim signal fed back to control the speed of the backward motion on the grating as described in (Ahrens et al., 2013).

## Computational Methods

### Lag-1 cross-correlation calculation for composite movie analysis

We calculated ⟨*XY*⟩_1_, the lag-1 cross-correlation as follows. First, we denote the cross-correlation function of *X* and *Y* as (*X* * *X*)(*t*), which is a discrete function centered at *t=0. t* denotes the lag. Then,

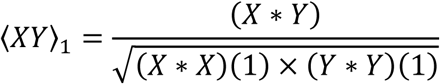

which is a function of time. The lag-1 cross-correlation reduces the effect of uncorrelated shot noise on the correlation between noisy signals *X* and *Y*.

### PCA-ICA analysis

All PCA-ICA analyses were conducted on full movies that had been denoised following SGPMD-NMF. The movies were then high-pass filtered in time with a 50 Hz high-pass filter. The high pass filtered movies were then segmented using PCA followed by time-domain ICA (Mukamel et al., 2009). The maximum number of sources from PCA-ICA was set to 20, and further components that did not correspond to cells were eliminated manually by inspection of the temporal and spatial components.

### SGPMD-NMF

Code to run SGPMD-NMF on an example dataset is available on GitHub here: https://github.com/adamcohenlab/invivo-imaging. Instructions for installing and running the code are here: http://bit.ly/sgpmdnmf-instructions. All SGPMD-NMF analyses done in the paper were run on the Harvard FASRC Cannon cluster.

## Acknowledgements

We thank C. Cai, A. Giovannuci, A. Singh, and K. Svoboda for Voltron data. We thank T. Kawashima, M. Ahrens, E. Jung, K. Piatkevich, and E. Boyden for zArchon1 zebrafish transgenics. The computations in this paper were run on the FASRC Cannon cluster supported by the FAS Division of Science Research Computing Group at Harvard University. This work was supported by the Harvard Data Science Initiative, by NIH grant R01MH117042 and by the Howard Hughes Medical Institute.

## Supplemental Figures

**Supplemental Figure S1.**
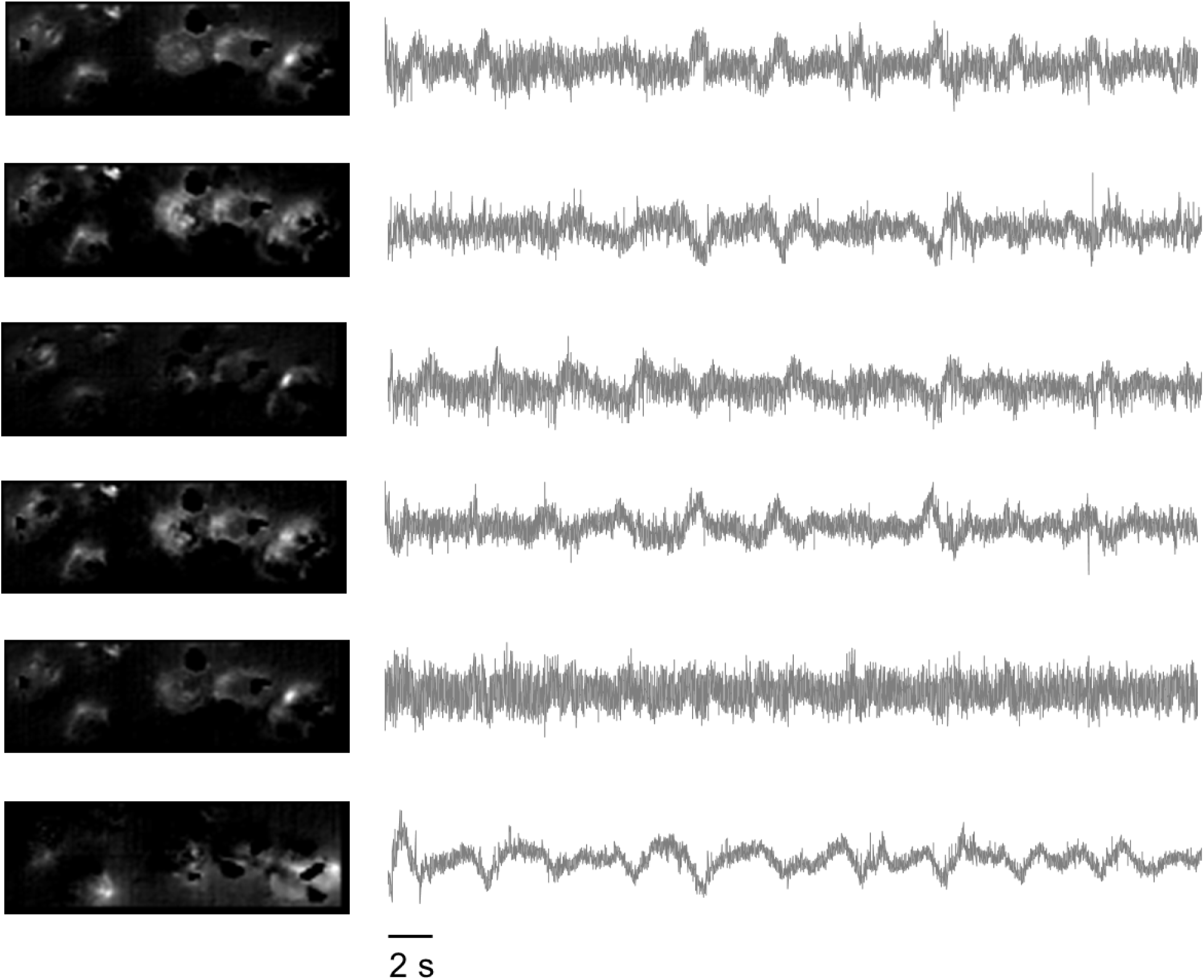
Background traces for mouse hippocampus CA1 SomArchon data. Spatial and temporal components of the background identified by SGPMD-NMF. Only background components that have a total variance across all pixels of at least two percent of the total variance in the denoised movie are shown.

**Supplemental Figure S2.**
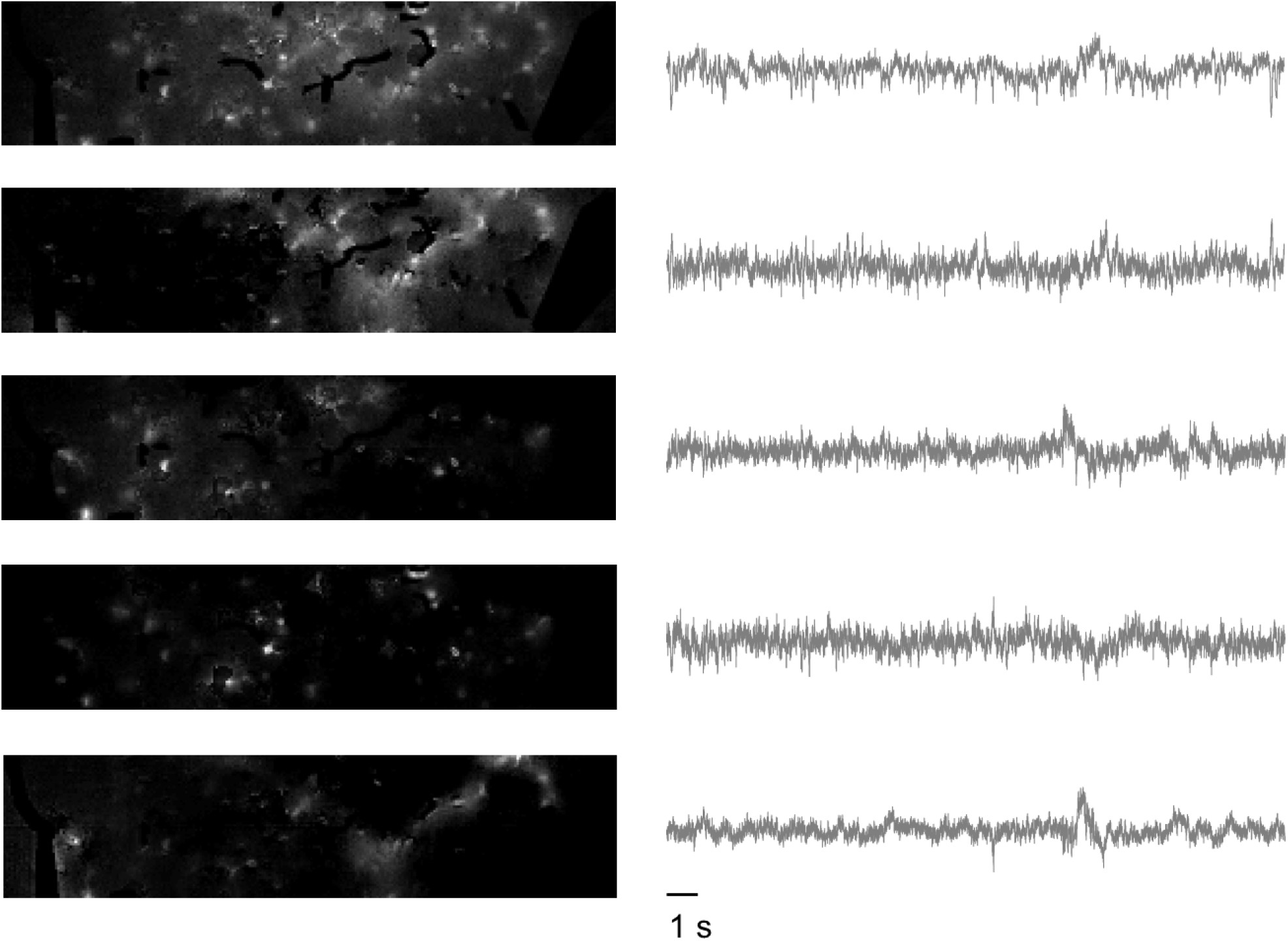
Background traces for mouse cortex L1 Voltron data. Spatial and temporal components of the background identified by SGPMD-NMF. Only background components that have a total variance across all pixels of at least two percent of the total variance in the denoised movie are shown.

**Supplemental Figure S3.**
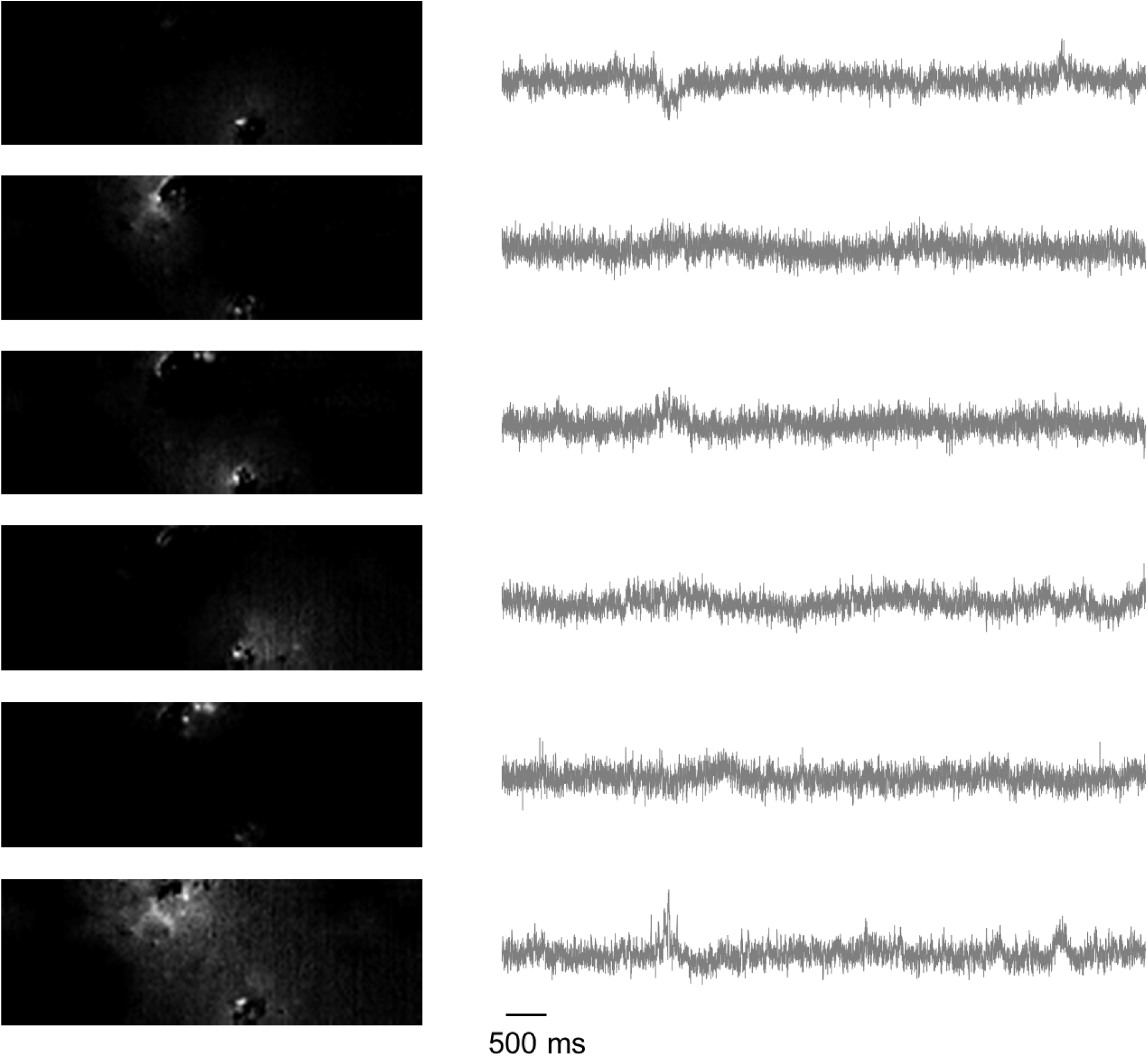
Background traces for mouse cortex L1 SomArchon data. Spatial and temporal components of the background identified by SGPMD-NMF. Only background components that have a total variance across all pixels of at least two percent of the total variance in the denoised movie are shown.

**Supplemental Figure S4.**
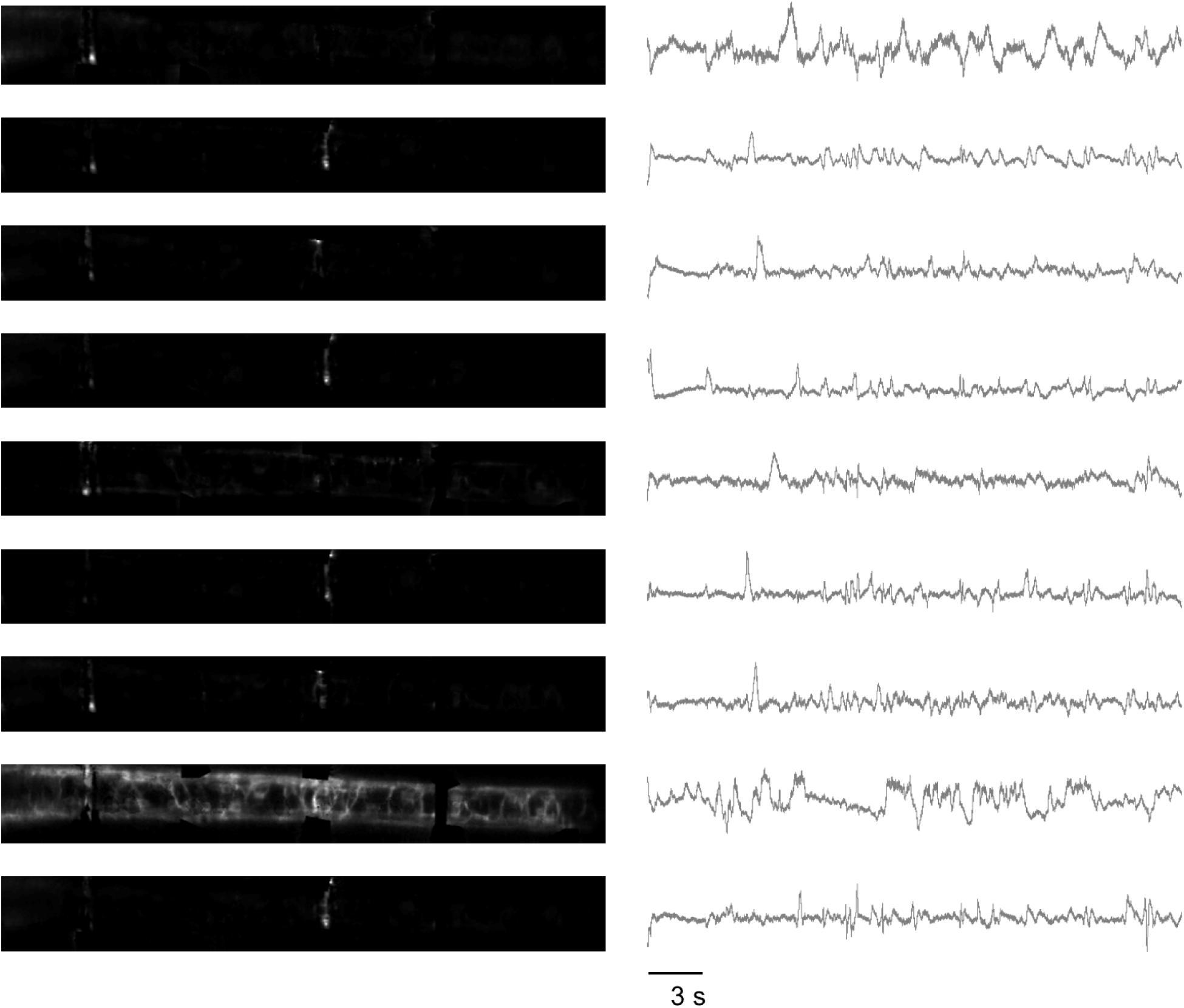
Background traces for zebrafish spine zArchon1 data. Spatial and temporal components of the background identified by SGPMD-NMF. Only background components that have a total variance across all pixels of at least two percent of the total variance in the denoised movie are shown.

## Notes

https://github.com/adamcohenlab/invivo-imaging

http://bit.ly/sgpmdnmf-instructions

https://drive.google.com/open?id=114nBlwpTjVEpxft65KBlJB4qfoczZM_p

https://drive.google.com/open?id=14W_Xzj3VzhC6lnpC86WyHEfC9_o33Wzm

https://drive.google.com/open?id=14GRnAe7Szl5R2NMwhtfXkPgfK9HSFUgS

https://drive.google.com/open?id=1qtCV1uyOmPl1Hjw3oGjt1xe5NtViQDKr

https://drive.google.com/open?id=1r9Vg3smCverw1B4O5yP5pabQrrURb-HP

